# Frozen Protein Foundation-Model Embeddings Improve Antibody–Antigen Binding Affinity Prediction

**DOI:** 10.64898/2026.07.13.738250

**Authors:** Roger Wang, Kevin Jin, Lurong Pan

## Abstract

We investigate whether representations from AINN-P1—a protein foundation model trained autoregressively on tens of millions of natural protein sequences—transfer to the task of ranking antibody–antigen pairs by binding affinity. Casting affinity maturation as a learning-to-rank problem over the change in binding free energy (ΔΔG), we compare a task-specific sequence model trained end-to-end from scratch against lightweight downstream heads built on top of frozen AINN-P1 embeddings, all evaluated under an identical five-fold cross-validation protocol. A regularized linear probe on the frozen embeddings already surpasses the from-scratch baseline, and an optimized lightweight head raises the mean Spearman rank correlation from 0.42 to 0.53—a relative improvement of approximately 28%— while training in seconds and without any fine-tuning of the foundation model. Because a linear probe alone exceeds the fully trained end-to-end baseline, the gain is attributable to representation quality rather than to added downstream-model capacity. These results position frozen foundation-model embeddings as a strong, data-efficient default for affinity ranking in antibody engineering and establish a conservative lower bound that task-adaptive fine-tuning is expected to exceed.

## 1. Introduction

Antibody affinity maturation requires ranking large panels of candidate variants by predicted binding affinity, often under stringent experimental-label budgets. Predictors trained from scratch for each new target tend to be data-hungry, slow to train, and difficult to reproduce across campaigns. Protein foundation models offer an appealing alternative: a single network, pretrained self-supervised on large sequence corpora, learns broadly transferable biochemical and evolutionary signal that can be reused across many downstream tasks (Alley et al., 2019; Rives et al., 2021; Lin et al., 2023).

Protein language models learn representations directly from unlabeled sequences. Autoregressive and masked models such as UniRep (Alley et al., 2019) and the ESM family (Rives et al., 2021; Lin et al., 2023) have shown that self-supervised pretraining on large corpora such as UniRef (Suzek et al., 2015) yields embeddings that transfer to diverse structural and functional tasks. Large-scale benchmarks such as ProteinGym (Notin et al., 2023) further show that, for protein fitness prediction, representation quality frequently dominates downstream-model complexity. AINN-P1 follows this paradigm, using an autoregressive next-token objective over tens of millions of natural sequences.

In this work we quantify the transfer value of AINN-P1 for antibody–antigen ΔΔG ranking. We treat the foundation model as a frozen encoder and train only a lightweight supervised head on top of its embeddings, and we contrast this against a task-specific model that learns its own representation end to end (Figure 1). Our contributions are threefold:

**Figure 1.**
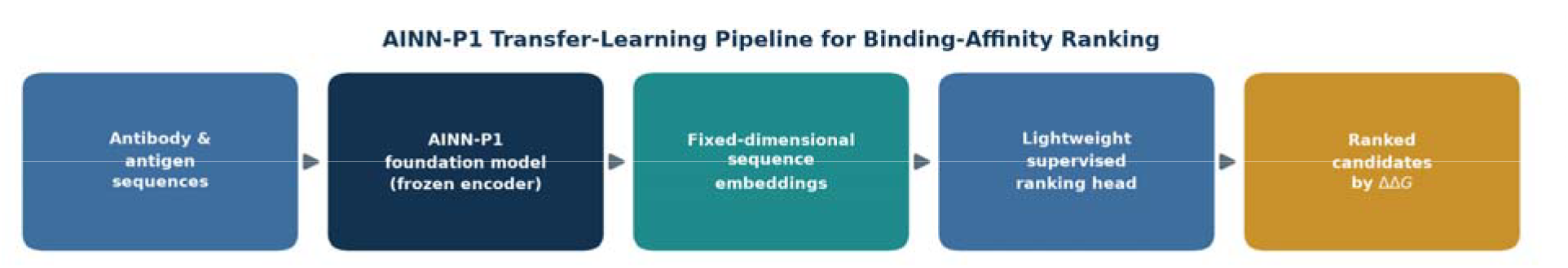
AINN-P1 transfer-learning pipeline. Antibody and antigen sequences are encoded by the frozen AINN-P1 foundation model; the resulting fixed-dimensional pair embedding feeds a lightweight supervised ranking head that orders candidates by predicted ΔΔG.

- A controlled comparison, under identical five-fold cross-validation, of frozen foundation-model embeddings versus a from-scratch baseline for ΔΔG ranking.
- Evidence that a linear probe on frozen AINN-P1 embeddings already exceeds the from-scratch model, isolating representation quality as the source of the gain.
- An optimized lightweight head that improves mean Spearman ρ from 0.42 to 0.53 (≈ 28% relative) at a small fraction of the training cost, with the foundation model held frozen.

## 2. Results

An overview of the transfer-learning pipeline is shown in Figure 1. Antibody and antigen sequences are encoded independently by the frozen AINN-P1 foundation model; the resulting fixed-dimensional pair embedding feeds a lightweight supervised ranking head that orders candidates by predicted ΔΔG.

We evaluated three configurations on a curated antibody–antigen ΔΔG dataset under a shared five-fold cross-validation protocol (Materials and Methods). All configurations used identical inputs, labels, and folds, so that differences in performance reflect only the representation and downstream head rather than the data split.

Frozen AINN-P1 embeddings improved ranking quality at every level of downstream-model complexity (Table 1, Figure 2). A regularized linear probe on the frozen embeddings already exceeded the from-scratch baseline, raising the mean five-fold Spearman correlation from ρ = 0.417 to ρ = 0.457. Replacing the linear probe with an optimized lightweight nonlinear head increased performance further to ρ = 0.533—an improvement of approximately 28% relative to the from-scratch baseline—while the foundation model remained frozen throughout.

**Table 1.** Mean five-fold cross-validated Spearman ρ for antibody–antigen ΔΔG ranking. All configurations use identical data and folds; the best configuration is shown in bold.

| Configuration | Representation | Mean 5-fold Spearman $\rho$ |
| --- | --- | --- |
| Task-specific model (from scratch) | Learned end-to-end | 0.417 |
| AINN-P1 embeddings + linear head | Frozen AINN-P1 | 0.457 |
| **AINN-P1 embeddings + optimized head** | **Frozen AINN-P1** | **0.533** |

**Figure 2.**
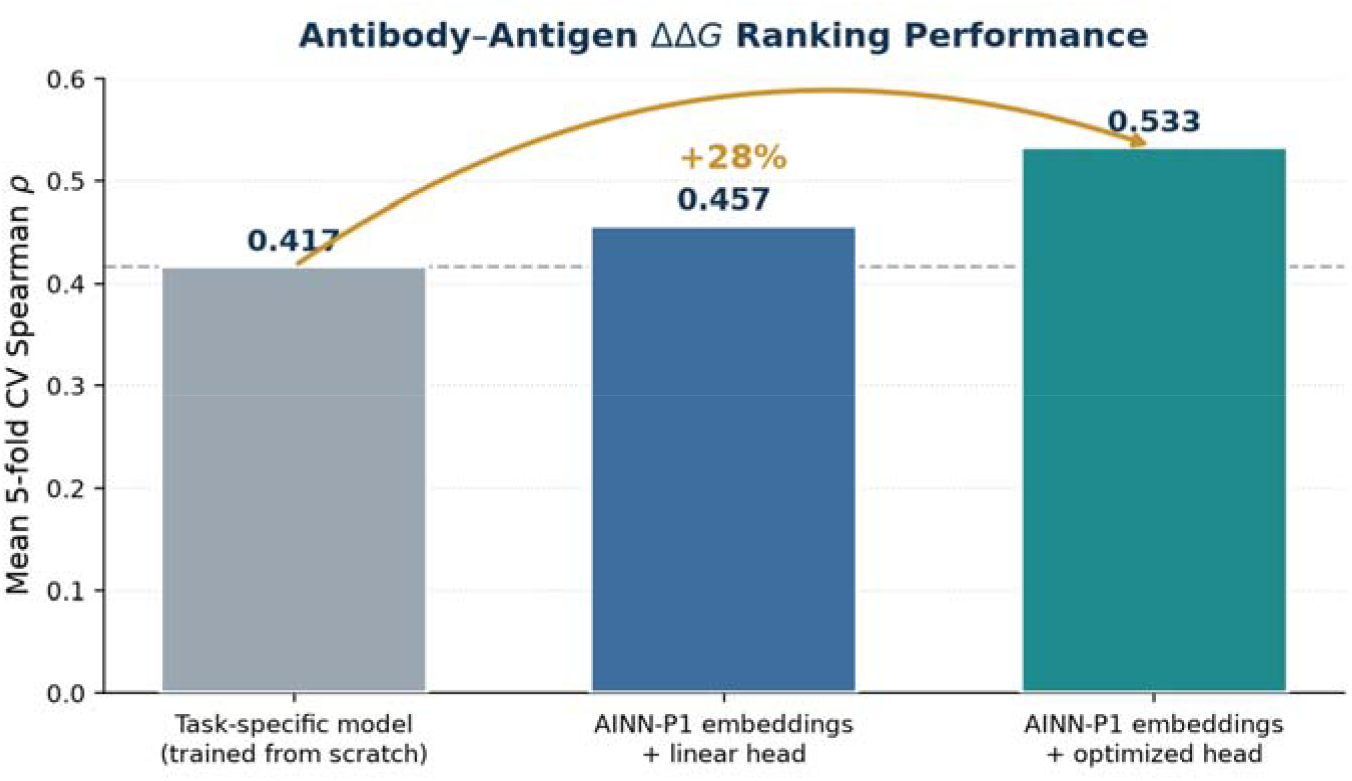
Ranking performance across configurations. Frozen AINN-P1 embeddings lift the mean five-fold Spearman ρ from 0.42 (from-scratch baseline) to 0.53 (optimized head), a ≈ 28% relative improvement.

Two features of this pattern are notable. First, the ordering tracks representation quality rather than model capacity: the largest single jump comes from replacing the end-to-end learned representation with frozen AINN-P1 embeddings, not from adding capacity to the downstream head. Second, the linear probe— arguably the lowest-capacity supervised model considered—already outperforms the fully trained end-to-end network, indicating that the frozen embeddings expose affinity-relevant structure in a nearly linearly separable form.

## 3. Discussion

The pattern of results points to representation quality as the primary driver of performance. Self-supervised pretraining encodes biochemical and evolutionary constraints that act as a strong inductive prior, so a small downstream head can generalize well from limited labels in a regime where an end-to-end model tends to overfit. The most direct evidence is that a linear probe on frozen embeddings already outperforms the from-scratch model: the gain originates in the representation, not in added downstream capacity.

Practically, the frozen-encoder design is modular, fast, data-efficient, and reproducible. The same embeddings can be reused across targets and downstream objectives, each downstream head trains in seconds rather than hours, and the absence of foundation-model fine-tuning removes a major source of run-to-run variability. Together these properties make frozen AINN-P1 embeddings a practical default for affinity ranking within iterative discovery and affinity-maturation campaigns.

Because the foundation model is held frozen, these results represent a conservative lower bound on achievable accuracy. A task-adaptive fine-tuning stage that specializes AINN-P1 to antibody–antigen affinity data is a natural next step and is expected to widen the margin further. Additional directions include higher-capacity AINN-P1 variants for richer embeddings, multi-objective downstream heads that jointly optimize affinity alongside developability or specificity, and closed-loop integration of experimental feedback. A current limitation is that evaluation is confined to a single curated ΔΔG dataset and to rank-based metrics; broader benchmarking across targets and prospective wet-lab validation are important next steps.

## 4. Materials and Methods

### 4.1 Task definition and metrics

We consider the antibody–antigen binding-affinity change, ΔΔG, which measures how a candidate’s binding free energy differs from a reference. Because downstream decisions—which candidates to advance to the wet lab—depend on the relative ordering of candidates rather than on absolute values, we frame the problem as learning to rank and evaluate it with rank-based metrics. The primary metric is the Spearman rank correlation (ρ) between predicted and measured ΔΔG, where higher ρ indicates better recovery of the true affinity ordering. We additionally track Normalized Discounted Cumulative Gain (NDCG) and area under the ROC curve (AUC) as complementary top-of-list and binary-separation measures.

### 4.2 Foundation-model representation

Each antibody and antigen sequence is encoded independently by AINN-P1 operating as a frozen encoder; no gradients propagate into the foundation model. AINN-P1 is a protein foundation model trained with an autoregressive next-token objective on tens of millions of natural protein sequences. The per-sequence embeddings are combined into a single fixed-dimensional representation of the antibody–antigen pair, which serves as the input to all downstream models (Figure 1).

### 4.3 Downstream ranking head

Because the frozen representation already captures rich sequence signal, only a compact supervised model is required to map it to relative binding affinity. We evaluated a small family of lightweight heads, ranging from a regularized linear probe to an optimized nonlinear estimator, and report both the linear probe (as a representation-quality lower bound) and the best-performing configuration.

### 4.4 From-scratch baseline

As a reference, we trained a task-specific sequence model that learns its own representation end to end, with no foundation-model pretraining. All approaches share identical inputs, labels, and cross-validation splits, isolating the effect of the representation. Architectural and training details of the internal models are proprietary and are omitted here.

### 4.5 Cross-validation and evaluation

All approaches were assessed with five-fold cross-validation on a curated antibody–antigen ΔΔG dataset, using the same folds throughout so that scores are directly comparable. Feature normalization was fit only on training folds to prevent information leakage, and all reported metrics were averaged across folds. Spearman ρ served as the primary reporting metric, with NDCG and AUC monitored as secondary measures.

## 5. Conclusion

Pairing frozen AINN-P1 foundation-model embeddings with a lightweight downstream head improved antibody–antigen ΔΔG ranking from a Spearman ρ of 0.42 to 0.53—approximately a 28% relative gain— under identical cross-validation and at a small fraction of the training cost. The result demonstrates the transfer value of protein foundation-model representations and provides a validated basis for fine-tuned, production-grade affinity predictors in antibody engineering.

## Declarations

### Author contributions

R.W. and K.J.: methodology, software, formal analysis, investigation, visualization, and writing – original draft. L.P.: conceptualization, supervision, resources, funding acquisition, and writing – review & editing. All authors read and approved the final manuscript.

### Data and code availability

The AINN-P1 model and the curated antibody–antigen ΔΔG dataset used in this study are proprietary to Ainnocence, Inc.; architectural and training details of the internal models are not disclosed. The aggregate results supporting the findings are reported in full in Table 1. Requests regarding data or model access may be directed to the corresponding author.

## Acknowledgments

The authors thank the Ainnocence research and engineering teams for infrastructure support and helpful discussions.

